# LSTM-PHV: Prediction of human-virus protein-protein interactions by LSTM with word2vec

**DOI:** 10.1101/2021.02.26.432975

**Authors:** Sho Tsukiyama, Md Mehedi Hasan, Satoshi Fujii, Hiroyuki Kurata

**Affiliations:** Department of Bioscience and Bioinformatics, Kyushu Institute of Technology, 680-4 Kawazu, Iizuka, Fukuoka 820-8502, Japan; Japan Society for the Promotion of Science, 5-3-1 Kojimachi, Chiyoda-ku, Tokyo 102-0083, Japan

**Keywords:** LSTM, word2vec, protein-protein interaction, human, virus, prediction, amino acid sequence, deep learning, SARS-CoV2

## Abstract

Viral infection involves a large number of protein-protein interactions (PPIs) between human and virus. The PPIs range from the initial binding of viral coat proteins to host membrane receptors to the hijacking of host transcription machinery. However, few interspecies PPIs have been identified, because experimental methods including mass spectrometry are time-consuming and expensive, and molecular dynamic simulation is limited only to the proteins whose 3D structures are solved. Sequence-based machine learning methods are expected to overcome these problems. We have first developed the LSTM model with word2vec to predict PPIs between human and virus, named LSTM-PHV, by using amino acid sequences alone. The LSTM-PHV effectively learnt the training data with a highly imbalanced ratio of positive to negative samples and achieved an AUC of 0.976 with an accuracy of 98.4% using 5-fold cross-validation. By using independent test dataset, we compared the LSTM-PHV with existing state-of-the-art PPI predictors including DeepViral. In predicting PPIs between human and unknown or new virus, the LSTM-PHV presented higher performance than the existing predictors when they were trained by multiple host protein-including datasets. LSTM-PHV learnt multiple host protein sequence contexts more efficiently than the DeepViral. Interestingly, learning of only sequence contexts as words presented remarkably high performances. Use of uniform manifold approximation and projection demonstrated that the LSTM-PHV clearly distinguished the positive PPI samples from the negative ones. We presented the LSTM-PHV online web server that is freely available at http://kurata35.bio.kyutech.ac.jp/.

## Introduction

Viral infections are one of the major causes of human health, as we can see from the current status of the SARS-CoV2 that raises a global pandemic. As of February 2021, more than 110 million people infected and nearly 2.4 million deaths have been reported worldwide for the COVID-19 disease [1]. Viruses achieve their own life cycle and proliferate their clones by hijacking and utilizing the functions of their hosts. In the process to achieve this purpose, viruses interact with host proteins to control cell cycles and apoptosis and to transport their own genetic material into the host nucleus [2, 3]. Therefore, it is important to identify human-virus protein-protein interactions (HV-PPIs) in understanding the mechanisms of viral infections and host immune responses and finding new drug targets. However, compared to intraspecies PPIs, few interspecies PPIs have been identified. In identifying interactions, experimental methods such as yeast-to-hybrid and mass spectrometry have been widely used, but they are time-consuming and laborious. For this reason, it is difficult to apply experimental methods for all protein pairs. Therefore, the computational approach is a preliminary treatment prior to the experimental method.

The use of amino acid sequence information is promising in the prediction of PPIs because the experimental data of PPIs and sequence information of proteins are abundant. Machine learning (ML)-based approaches are very attractive [4] that use the amino acid binary profiles [5, 6], evolutionary properties [7, 8], physicochemical properties [9, 10], and structural information [11]. Zhou et al integrated different encoding methods, such as relative frequency of amino acid triplets, frequency difference of amino acid triplets, and amino acid composition to construct a SVM-based PPI predictor [12]. Recently, promising encoding schemes have been proposed to capture the sequence patterns of proteins, including conjoint triad [13, 14], auto covariance [15], and autocorrelation [16].

Human-virus interactions involve not only the various properties of amino acid sequences but also the arrangement of 20 amino acid residues in the semantic context of whole protein sequences. While many predictors have focused on the former features, the latter context-based information is suggested to be effective in predicting HV-PPIs [17]. To capture the context information of amino acid residue sequences as much as possible, word/document embedding techniques have recently been proposed. Yang et al. combined the doc2vec encoding schemes with a random forest method to predict PPIs [17]. The DeepViral (Liu-Wei, et al., 2020) combined doc2vec/word2vec embedding methods with a convolutional neural network (CNN). DeepViral also encoded host phenotype associations from PathoPhenoDB [18] and protein functions from the Gene Ontology (GO) database (The Gene Ontology Consortium, 2017) to predict PPIs. In addition, several constructed ML models were designed for certain individual virus species, limiting their generalizability to other human host-virus systems [19-21].

To utilize the amino acid sequence context as words effectively, we have proposed the long short-term memory (LSTM) model [22] with the word2vec embedding method that predicts the PPIs between human and virus, named LSTM-PHV. To the best of our knowledge, this is the first application of the LSTM with the word2vec to sequence-based PPI prediction. Interestingly, use of the sequence context as words presented remarkably accurate prediction of the interactions between human and unknown virus proteins.

## Materials and methods

### Datasets construction

The data of PPIs were downloaded from the Host-Pathogen Interaction Database 3.0 (HPIDB 3.0) [23]. The retrieved HV-PPIs were further selected by the following process. First, to ensure interactions with a certain level of confidence, the PPIs with an MI score of below 0.3 were removed. The MI score is the confidence score assigned to each PPI from IntAct [24] and VirHostNet [25]. Second, redundant PPIs were excluded by using CD-HIT with an identity threshold of 0.95 [26]. Third, the PPIs that contained the proteins consisting of standard amino acids only and the proteins with a length of more than 30 residues and less than 1000 residues were selected. Finally, 22383 PPIs from 5882 human and 996 virus proteins were considered as positive samples.

To the best of our knowledge, there is no gold standard for generating negative samples. Many previous studies used a random sampling method. Pairs of the human and virus proteins that do not appear in the positive PPI dataset are randomly sampled as negative data. However, the random sampling method may incorrectly assign many positive samples to negative ones [5, 20]. To address this problem, the dissimilarity negative sampling method was developed [5], which used a sequence similarity-based method to explore the protein pairs that are unlikely to interact. We employed the dissimilarity-based negative sampling method as follows. We calculated the sequence similarities of all pairs of virus proteins in positive samples with the Needleman-Wunsch algorithm of BLOSUM30 and defined a similarity vector for each virus protein. Subsequently, we excluded the virus proteins showing lower sequence similarities than *Ts* for more than half of the total virus proteins as outliers. *Ts* was calculated by:

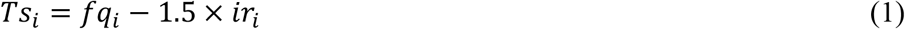

where *fq*_*i*_ and *ir*_*i*_ are the first quartile and quartile range of the similarity scores for the *i*-th virus protein *V*_*i*_, respectively. By setting the maximum and minimum values of the similarity scores to 0 and 1, respectively, the similarity score was normalized and converted into the distance.

The human proteins that consisted of the standard amino acids and whose residue length was longer than 30 and shorter than 1000 were retrieved from the SwissProt database [27]. Then, the human proteins that interacted with the virus proteins showing a distance from viral protein *V*_*i*_ of less than distance threshold *T* were removed, considering that they were likely to interact with virus protein *V*_*i*_. The remaining pairs of the human and virus proteins were regarded as negative samples. According to the previous study [5], the threshold *T* was set to 0.8. We randomly sampled from the candidates so that the ratio of positive to negative samples was 1:10. The positive and negative samples were labeled 1 and 0, respectively. The resultant dataset was divided into training data and independent test data at a ratio of 8:2.

To assess whether a predictor is applicable to unknown or new virus species, we employed the four training datasets (TR1: PPIs between human and any virus except Influenza A virus subtype H1N1 (H1N1), TR2: PPIs between human and any virus except *Ebola virus*, TR3: PPIs between any host and any virus except H1N1, TR4: PPIs between any host and any virus except *Ebola virus*) and two test datasets (TS1: PPIs between human and H1N1 virus; TS2: PPIs between human and Ebola virus). They were provided by Zhou et al [12].

### Embedding of protein sequences by word2vec

In the field of natural language processing, embedding methods such as word2vec [28] and doc2vec [29] were developed to obtain the distributed representation of words and documents, respectively. In word2vec, the weights in a neural network learn the context of words to provide the distributed representation that encodes different linguistic regularities and patterns [30]. There are two methods for learning the context of words: Continuous Bag-of-Words Model (CBOW) and the Continuous Skip-Gram Model (Skip-Gram). CBOW predicts the current word based on the context, while Skip-Gram predicts the context from the current word. Skip-gram is more efficient with less training data, while CBOW learns faster and more frequent words. At present, computational biology used these methods [31, 32].

The amino acid sequences of human and virus proteins registered as positive and negative samples were encoded as matrixes using the word2vec method. The k-mers (k consecutive amino acids) in amino acid sequences were regarded as a single word (unit) and each amino acid sequence was represented by multiple k-mers. For example, given an amino acid sequence MAEDDPYL, the unit of the 4-mers are MAED, AEDD, EDDP, DDPY, and DPYL (Fig. 1). We trained a CBOW-based word2vec model to learn the appearance pattern of k-mers from the computational speed standpoint by using the genism of the python package [33]. Here, k-mers and protein sequences correspond to words and sentences in natural language. Human and virus proteins in positive samples and non-redundant proteins in the SwissProt database [27] were used to train the word2vec model. The non-redundant proteins were collected by applying CD-HIT to all proteins with an identity threshold of 0.9. The k-mers up to three neighbors of a specific k-mer are considered as the peripheral k-mers, and training was iterated 1000 times. The trained word2vec model produced 128-dimensional embedding vectors in each k-mer and they were concatenated to produce the embedding matrixes of proteins. Since 4-mer provided the largest AUC by 5-fold cross-validation in a previous study [17], we set k to 4.

**Fig. 1.**
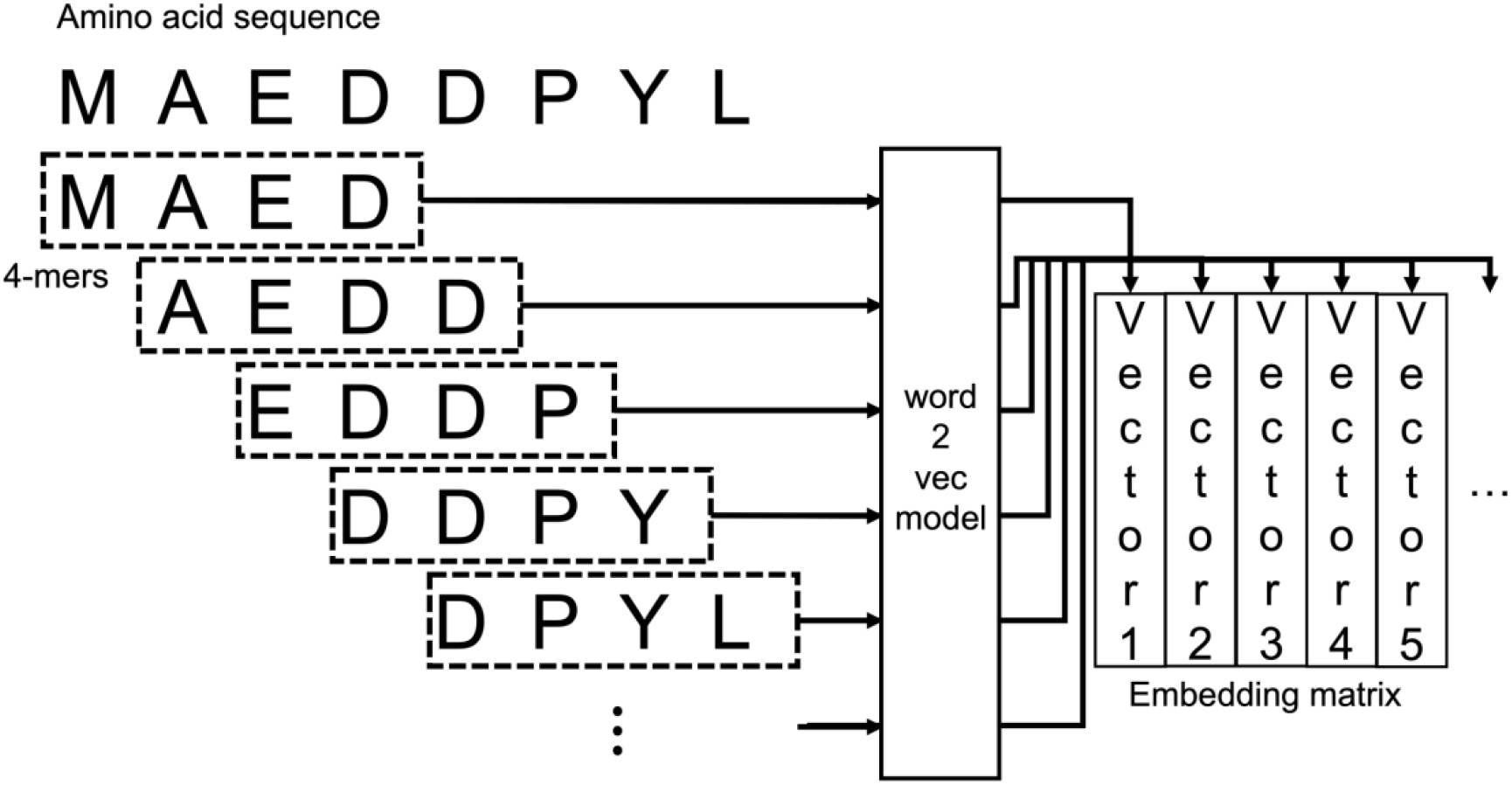
Embedding of amino acid sequences in a case of 4-mer. Amino acid sequences were represented by 4-mers and embedded as a matrix by training the word2vec model. The matrixes were generated by concatenating the vectors of 4-mers in a row.

### Construction of LSTM-PHV

Neural networks such as convolutional neural network (CNN) and recurrent neural network (RNN), in particular, are very powerful and have been applied to difficult problems such as speech recognition and visual object recognition [34]. The RNN learns time or step dependencies in sequence data and enables training on variable-length data. The LSTM solves the gradient explosion and gradient disappearance problems of RNNs, enabling long-term time-dependent learning.

The LSTM-PHV is composed of three sub-networks (Fig. 2). The two, upstream networks with the same structure transformed the human and virus proteins-embedding matrixes into two fixed-length vectors. The third network used their concatenated fixed-length vectors to predict the PPIs. They are referred to as “concatenated vectors”. The amino acid sequence column vectors in the embedding matrixes are inputted to each step of the LSTM units. The LSTM units were expanded in both the N-to C-terminus and the C-to N-terminus direction. The 64×2-dimensional vectors generated from one LSTM unit were concatenated in a row. The dimensions of the vectors generated through the three layers decreased in the order of 64, 32, and 1. In the first two layers, the rectified linear unit (ReLU) function with a dropout rate of 0.3 was used as an activation function. The scalar values generated from the third layer were lined up into a vector, which was provided to the softmax function. A fixed-length vector was generated by summarizing the weighted vectors in all the steps. To create a ML model for HV-PPI prediction, the key step is to conduct the feature encoding that converts human and virus protein sequences to the fixed-dimensional vectors.

**Fig. 2.**
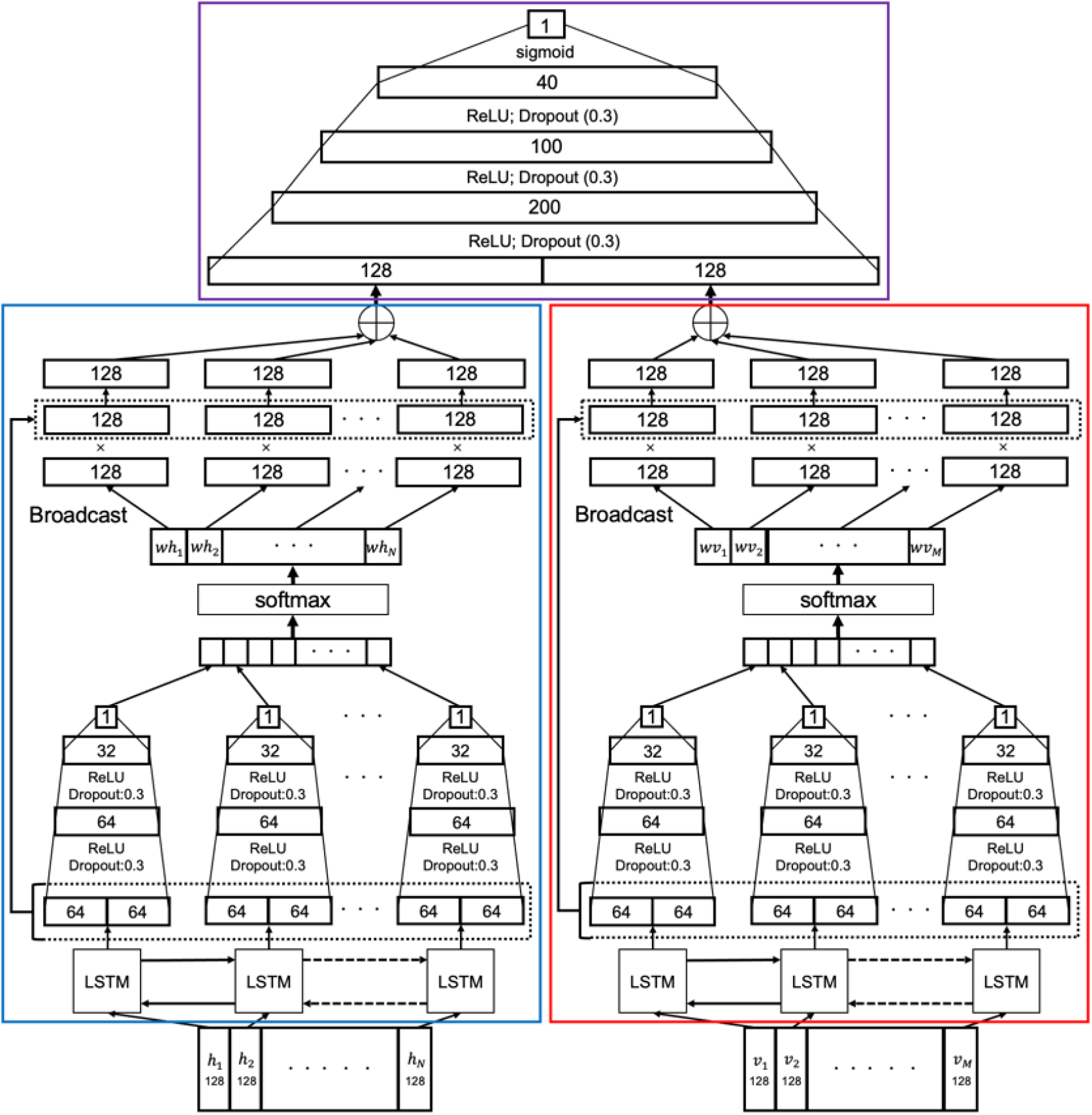
Three sub-neural networks of LSTM-PHV. The blue and red lines show the two, upstream neural networks with the LSTM that transform the human and virus protein matrices into the two fixed-length vectors, respectively. The purple line shows the final neural network that concatenates the fixed-length vectors of human and virus proteins to predict their PPIs.

The fixed-length vectors for the human and virus proteins were concatenated in line and propagated into the final network. The final network consists of three layers and an output layer. The dimensions of the generated vectors from each layer decreased in the order of 200, 100, 40, and 1. The ReLU function with a dropout rate of 0.3 was applied to the output of the three layers. To obtain a final output with a value between 0 and 1, the sigmoid function was used as an activation function at the output layer. The construction and learning of the neural networks were performed using the PyTorch [35] of the python package.

### Training of imbalanced data

Five-fold cross-validation was applied on the training dataset, while conserving the ratio of positive and negative samples at each subset. We set the learning rate to 0.001, used the rectified adam (RAdam) optimizer [36] as the optimization function, and set a mini-batch learning size to 1024. To train the model on imbalanced data, we weighted a binary cross-entropy loss function in the manner reported by Cui et al (Cui et al., 2019). The loss functions used are shown below.

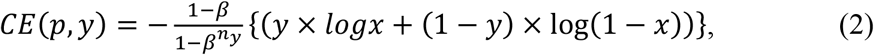

where *y* is the correct label, *x* is the model-predicted probability of interaction, *n*_*y*_ is the number of data whose label is *y* in the mini-batch, and *β* is the hyperparameter. *β* was set to 0.99. To prevent over-learning, the training process was terminated when the maximum accuracy in the validation data was not updated for consecutive 20 epochs. To prevent the weight of the loss function from being 0, we set an approximately equal ratio of labels for all the mini-batches.

### Measures

To evaluate the prediction performance, 7 statistics measures were used: sensitivity (recall), specificity, accuracy, Matthews correlation coefficient (MCC), precision, F1-score, area under the curve (AUC), and area under the precision-recall curve (AUPRC). MCC, F1-score, and AUPRC are effective in assessments of imbalanced data. The measures other than the AUC and AUPRC are given by:

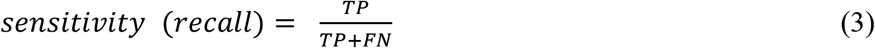

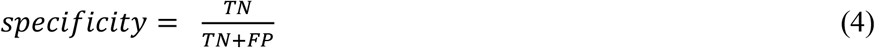

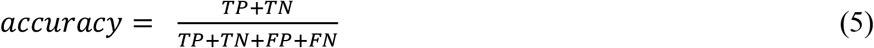

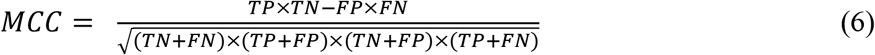

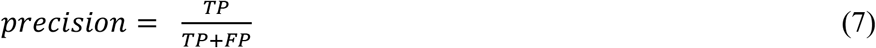

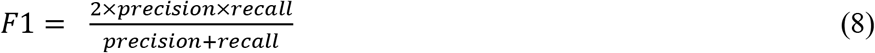

where TP, FP, TN and, FN are the numbers of the correctly predicted positive samples, incorrectly predicted positive samples, correctly predicted negative samples and incorrectly predicted negative samples, respectively. The threshold for a determination of whether protein pairs interact or not was set to a predicted probability of 0.5. AUC and AUPRC are the areas beneath the ROC curve and PR curve, respectively. These measures were calculated by the scikit-learn of the python package [37].

### Visualization of positive and negative samples

To visualize the concatenated vectors, we reduced the dimensionality of the concatenated vector from 256 to 2 using the uniform manifold approximation and projection (UMAP). UMAP is the nonlinear dimensionality reduction approach [38], which can preserve not only local patterns but also global patterns in low-dimensional space. In this study, the number of neighbors in the k-neighbor graph was set to 50, and a minimum distance between points in the low-dimensional space was set to 0. The distances between any points were calculated by the Euclidean distance. The optimization was implemented up to 500 epochs with a learning rate of 1.0.

## Results and Discussion

### Performance of LSTM-PHV

We used the LSTM-PHV to predict HV-PPIs using amino acid sequences alone. Prediction performances were evaluated via 5-fold cross-validation on the training dataset. Out of the five models, the model with the highest AUC was used to predict the independent test dataset. The accuracies on the training and independent datasets were 0.984 and 0.985, respectively (Fig. 3 and Fig. 4). The AUCs were 0.976 and 0.973 on the training and independent datasets, respectively (Tables S1 and S2). To characterize the performance of LSTM-PHV, we compared it with Yang’s model (RF model with Doc2vec) [17] on our independent test data, as shown in Fig. 4 and Table S2. The source code and trained model were provided by Yang et al with their recommended three threshold values. As in our case, Yang et al. used the imbalanced data that contained 10 times more negative samples than positive samples, generating negative samples by the dissimilarity-based negative sampling method. The LSTM-PHV presented higher values than Yang’s model not only for AUC and accuracy but also for MCC, F1-score, and AUPRC. The LSTM-PHV was able to learn the imbalanced data better than Yang’s model. The high MCC takes a great advantage, because learning of imbalanced data is essential. At present the number of known PPIs is very small compared to the total number of protein pairs. It is not evitable that negative samples are typically produced much more than positive ones in the absence of golden standard of generating negative samples.

**Fig. 3.**
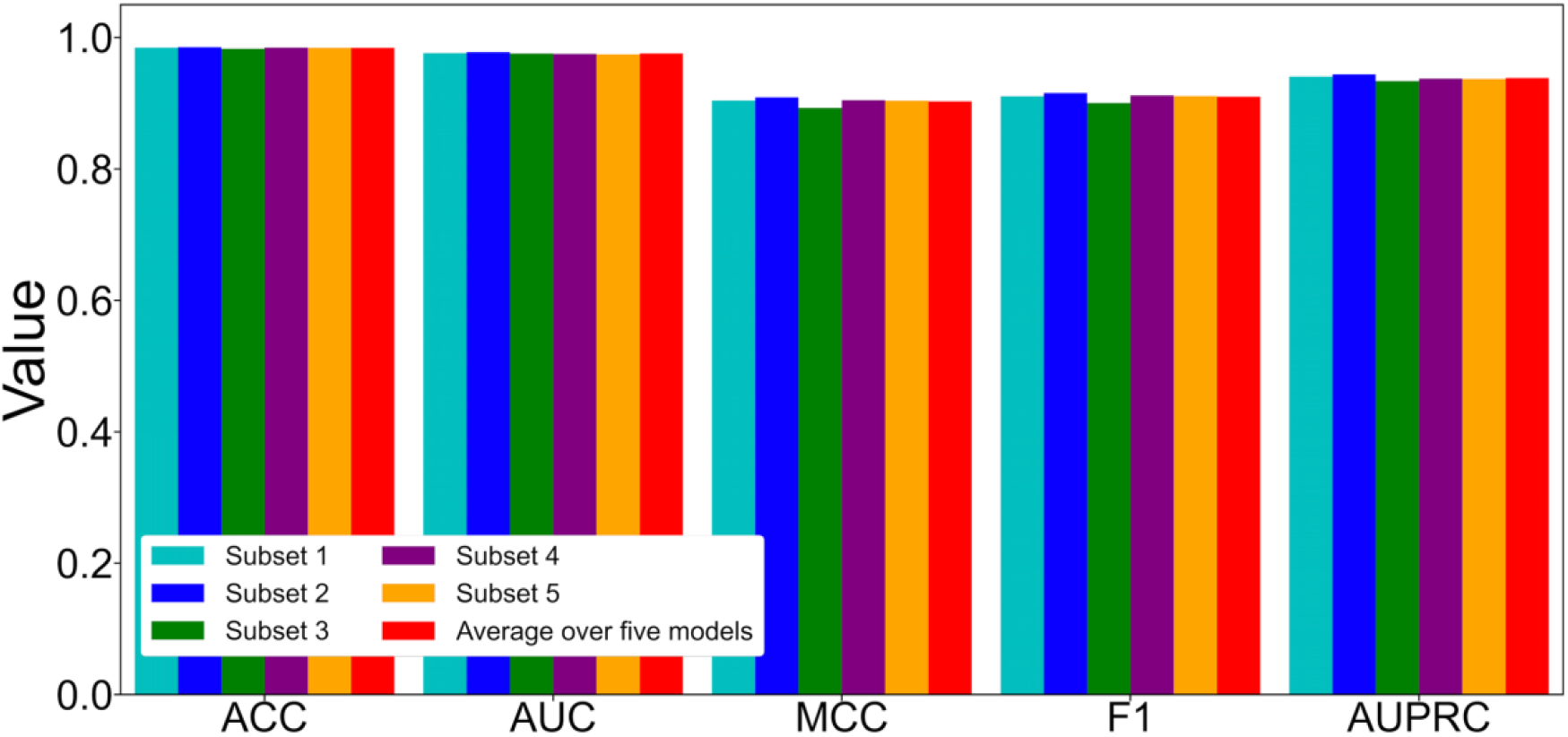
Performance of the LSTM-PHV by 5-fold cross validation on our training dataset. SN, SP, ACC, AUC, MCC, F1, and AUPRC correspond to sensitivity, specificity, accuracy, the area under the ROC curve, Matthews correlation coefficient, F1-score, and area under the Precision-Recall curve, respectively.

**Fig. 4.**
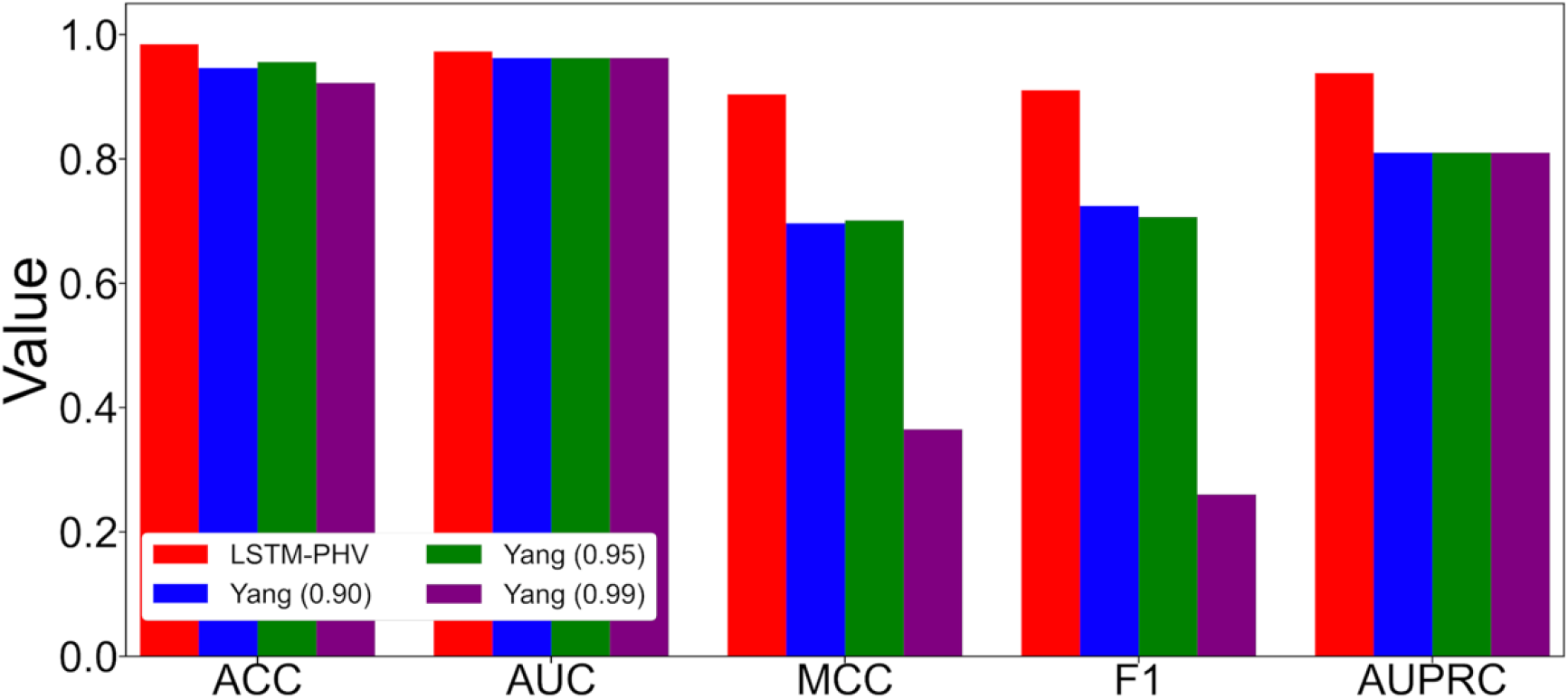
Performance comparison of LSTM-PHV with Yang’s RF model with doc2vec using our independent test. Thresholds at specificity of 0.90, 0.95 and 0.99 in Yang’s model are used according to their suggestion. SN, SP, ACC, AUC, MCC, F1, AUPRC correspond to sensitivity, specificity, accuracy, Matthews correlation coefficient, the area under the ROC curve, F1-score, area under the Precision-Recall curve, respectively.

### Predictive performance of new virus species

To assess whether the LSTM-PHV is applicable to unknown virus species, the LSTM-PHV was evaluated by using the four datasets provided by Zhou et al [12] (Fig. S2), which were employed by the DeepViral [39]. We compared LSTM-PHV with Zhou’s model and the DeepViral. In training the LSTM-PHV by Zhou’s datasets, we set a batch size to 256 and used the normal binary cross-entropy loss function, because Zhou’s datasets were much smaller than our dataset and it was balanced data. As shown in Fig 5 and Table S3, the LSTM-PHV presented higher performances than Zhou’s model (a SVM with commonly-used encoding methods) for all the datasets. The DeepViral used not only amino acid sequence contexts (as words) but also the two biological features, phenotype associations for viruses from PathoPhenoDB [18] and protein functions from the Gene Ontology (GO) database [40, 41]. We trained the LSTM-PHV and DeepViral predictors by the same datasets of TR1 and TR2 that do not include the PPIs between human and H1N1 and *Ebola virus*, respectively. The LSTM-PHV showed comparative prediction performance to the DeepViral, when only the sequence contexts were employed. Compared to the DeepViral that considered the biological features, the LSTM-PHV predicted the PPIs between human and H1N1 (TS1) and those between human and *Ebola virus* (TS2) with a little low accuracy and AUC. However, when the predictors were trained on multiple host protein-including TR3 and TR4 and the embedding features, which were associated with host and several virus species, were not available, the LSTM-PHV predicted the PPIs between human and H1N1 (TS1) with higher accuracy and AUC than the DeepViral and predicted the PPIs between human and *Ebola virus* (TS2) with higher accuracy. LSTM-PHV learnt multiple host protein sequence contexts more efficiently than the DeepViral. Interestingly, we revealed that learning of only sequence contexts as words presented remarkably high performances.

**Fig. 5.**
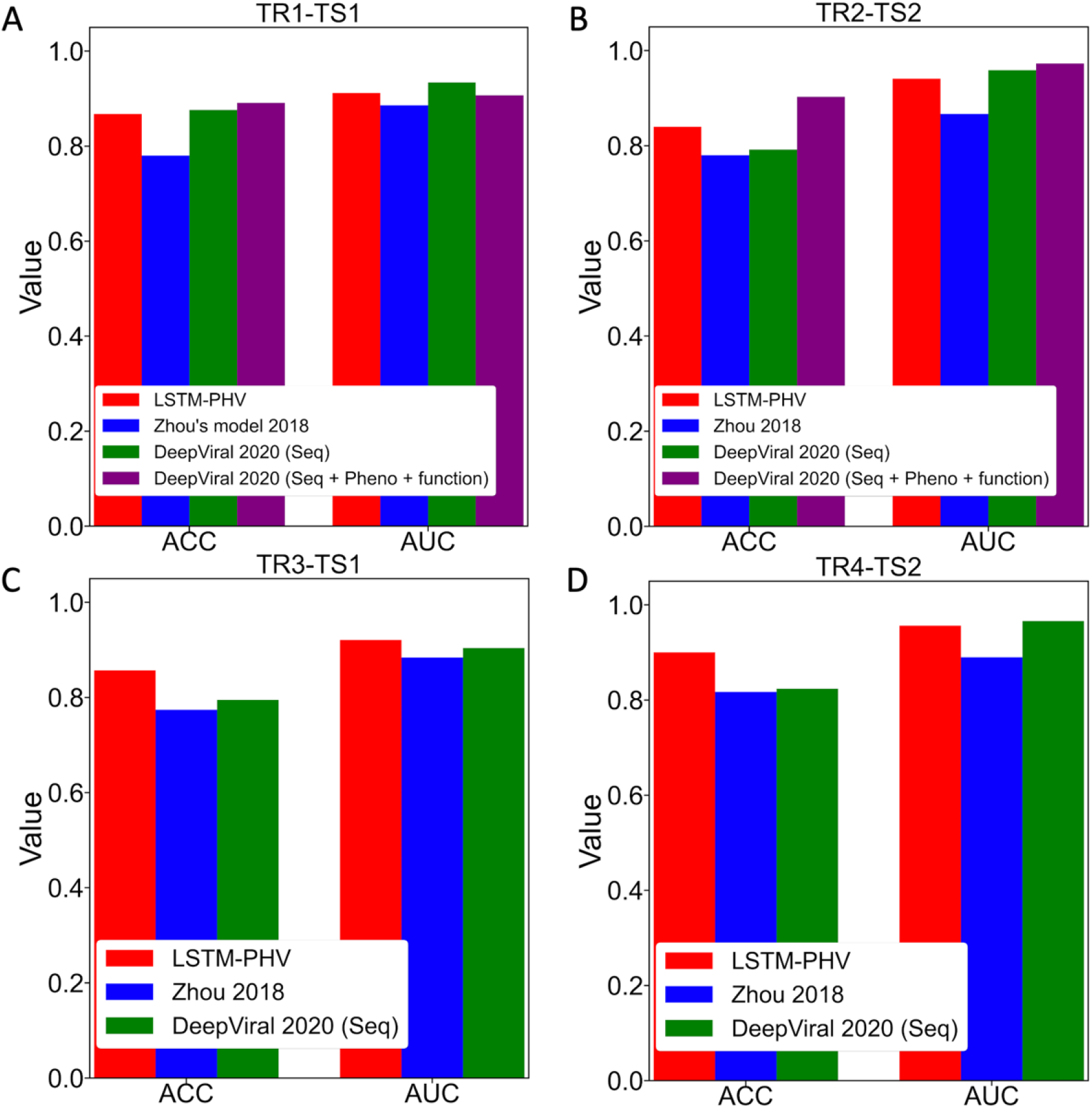
Prediction performance of LSTM-PHV with existing state-of-the-art predictors. We employed the four datasets (a, b, c, d) that combine the four training data with two test data according to Zhou’s study. The two datasets containing human-virus interactions (TR1-TS1 and TR2-TS2) were applied to LSTM-PHV, Zhou’s model, the DeepViral that used amino acid sequences alone, and the DeepViral integrating three features. The other two datasets containing host-virus interactions (TR3-TS1 and TR4-TS2) were applied to the LSTM-PHV, Zhou’s model and DeepViral. ACC and AUC correspond to accuracy and the area under the ROC curve, respectively. The performances were obtained from Table 1 in Liu-Wei’s paper (DeepViral). a) Performances in prediction of TS1 with the model trained on TR1 b) Performances in prediction of TS2 with the model trained on TR2 c) Performances in prediction of TS1 with the model trained on TR3 d) Performances in prediction of TS2 with the model trained on TR4

### Analysis of the vectors generated in deep neural networks

The two, upstream neural networks with the LSTM generated the fixed-length vectors. To examine how these neural networks extract PPI-related information, transforming the embedding matrices of proteins to the two fixed-length vectors, we drew the UMAP map of their concatenated vectors on the independent test data (Fig. 6 and Fig. S1-S4). In all the UMAP maps, multiple clusters were generated and positive samples were distinguished from negative samples within several clusters. The false negative and false positive samples were located between the true negative and true positive samples. The numbers of the false negative and false positive samples were small. These results suggested that the upstream neural networks extract critical information responsible for predicting PPIs from the amino acid sequences of each protein.

**Fig. 6.**
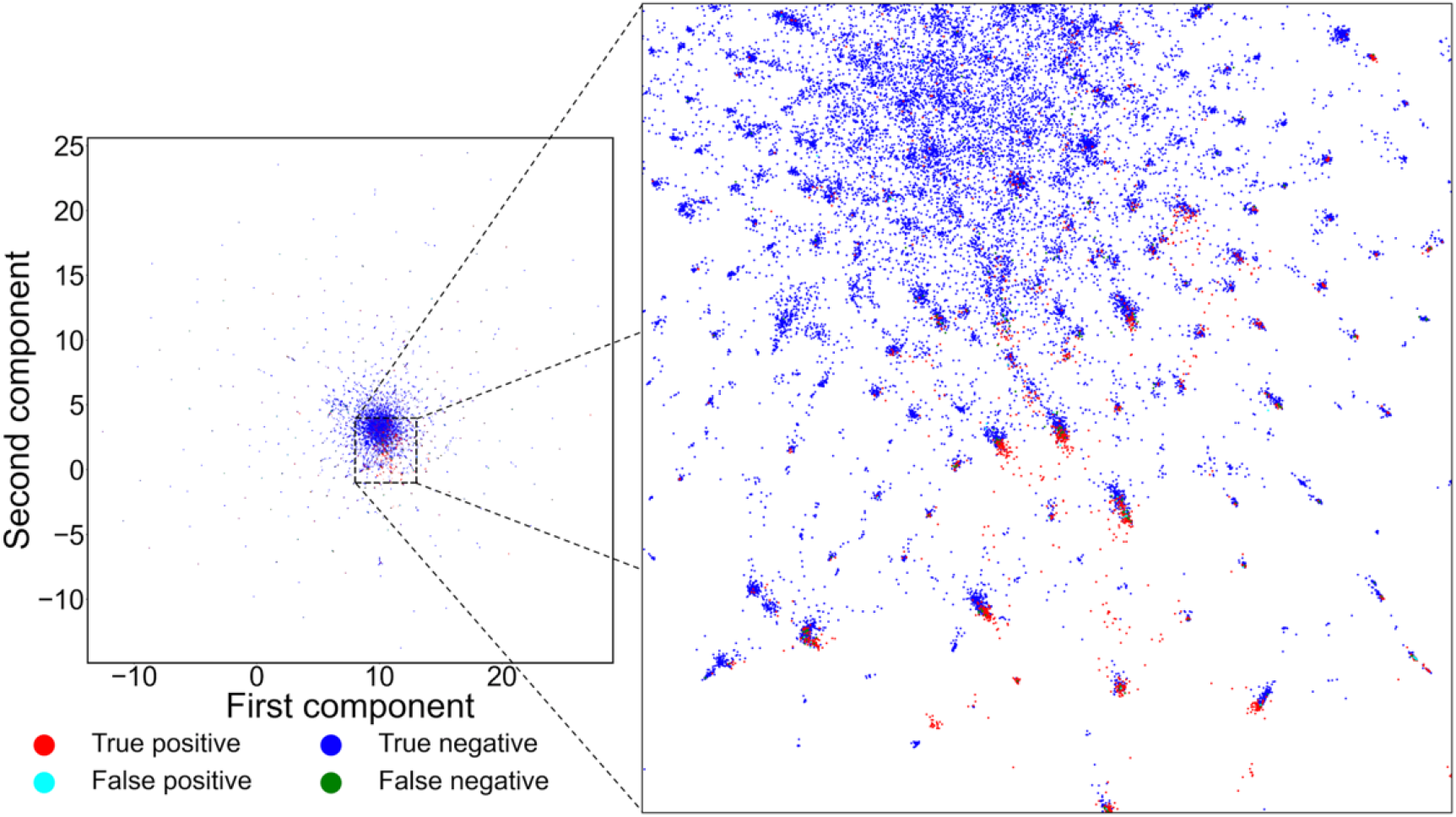
UMAP map of our independent test data. Many clusters were scattered. The true positive, false positive, false negative, and true negative samples were included by several clusters. True positive samples were distinguished from the true negative samples. The false negative and false positive samples were located between the true positive and true negative samples.

## Conclusions

We presented the LSTM-PHV that combined LSTM with the word2vec embedding method by considering the sequential amino acid arrangement alone to predict PPIs between human and virus. The LSTM-PHV consists of three neural networks: the two, upstream neural networks implement the LSTM units to transform the variable-length embedding matrices for human and virus proteins into the two fixed-length vectors, and the final neural network concatenates these fixed-length vectors to predict PPIs. The LSTM-PHV learnt highly imbalanced data and was able to accurately predict the interaction of a human protein to an unknown virus protein, compared to existing state-of-the-art models. The UMAP map of the concatenated vectors demonstrated that the LSTM-PHV clearly separated the positive samples from negative ones. Use of the LSTM-PHV enhances the screening of drug targets that inhibit human-virus PPIs and definitely contributes to advances in remedies of infectious diseases including COVID-19.

## Acknowledgement

This work was supported by the Grant-in-Aid for Scientific Research (B) (19H04208) and partially supported by the Grant-in-Aid for JSPS Research Fellow (19F19377) from Japan Society for the Promotion of Science (JSPS).

## Key points

The LSTM-based model with word2vec (LSTM-PHV) efficiently learnt highly imbalanced training data to accurately predict PPIs between human and virus.

Learning of amino acid sequence contexts as words are dominantly effective in prediction PPIs.

UMAP visualizes that positive samples are clearly distinguished from negative samples.

## Biographical note

**Sho Tsukiyama** is a student of Department of Interdisciplinary Informatics in the Kyushu Institute of Technology, Japan. His main research interests include machine learning and computational biology.

**Md Mehedi Hasan** received his PhD degree in bioinformatics from CAU, Beijing in 2016. He is currently as a JSPS international PD fellow in the Kyushu Institute of Technology, Japan. Before his current position, he worked as a researcher at Chinese University of Hong Kong, Hong Kong. His main research interests include protein structure prediction, machine learning, data mining, computational biology, and functional genomics.

**Satoshi Fujii** is an assistant professor of Department of Bioscience and Bioinformatics in the Kyushu Institute of Technology, Japan. His research interests include machine learning, clinical data analysis, and biomedical design.

**Hiroyuki Kurata** is a professor of Department of Bioscience and Bioinformatics in the Kyushu Institute of Technology, Japan. His research interests primarily focus on systems biology, synthetics biology, functional genomics, machine learning, and their applications.

