## Supplemental information for "LSTM-PHV: Prediction of human-virus protein-protein interactions by LSTM with word2vec"

**Table S1.** Performance of the LSTM-PHV by 5-fold cross validation on our training dataset.

| | SN | SP | ACC | AUC | MCC | PPV | F1 | AUPRC |
|---|---|---|---|---|---|---|---|---|
| **Subset 1** | 0.862 | 0.997 | 0.985 | 0.976 | 0.904 | 0.965 | 0.911 | 0.941 |
| **Subset 2** | 0.871 | 0.997 | 0.985 | 0.978 | 0.909 | 0.965 | 0.916 | 0.944 |
| **Subset 3** | 0.850 | 0.996 | 0.983 | 0.976 | 0.893 | 0.957 | 0.900 | 0.934 |
| **Subset 4** | 0.874 | 0.996 | 0.985 | 0.975 | 0.905 | 0.953 | 0.912 | 0.937 |
| **Subset 5** | 0.869 | 0.996 | 0.985 | 0.974 | 0.904 | 0.957 | 0.911 | 0.937 |
| **Average over five models** | 0.865 | 0.996 | 0.984 | 0.976 | 0.903 | 0.960 | 0.910 | 0.939 |

SN, SP, ACC, AUC, MCC, PPV, F1, and AUPRC correspond to sensitivity, specificity, accuracy, the area under the ROC curve, Matthews correlation coefficient, positive predictive value (precision), F1-score, and area under the Precision-Recall curve, respectively.

**Table S2.** Performance comparison of LSTM-PHV with Yang's RF model with doc2vec using our independent test.

| | SN | SP | ACC | AUC | MCC | PPV | F1 | AUPRC |
|---|---|---|---|---|---|---|---|---|
| **LSTM-PHV** | 0.862 | 0.997 | 0.985 | 0.973 | 0.904 | 0.965 | 0.911 | 0.938 |
| **Yang (0.90)** | 0.773 | 0.964 | 0.947 | 0.963 | 0.697 | 0.682 | 0.724 | 0.810 |
| **Yang (0.95)** | 0.587 | 0.993 | 0.956 | 0.963 | 0.701 | 0.888 | 0.707 | 0.810 |
| **Yang (0.99)** | 0.151 | 1.000 | 0.922 | 0.963 | 0.365 | 0.968 | 0.261 | 0.810 |

SN, SP, ACC, AUC, MCC, PPV, F1, AUPRC correspond to sensitivity, specificity, accuracy, the area under the ROC curve, Matthews correlation coefficient, positive predictive value (precision), F1-score, area under the Precision-Recall curve, respectively.

**Table S3.** Prediction performance of LSTM-PHV with existing state-of-the-art predictors.

| | | **LSTM-PHV** | **Zhou's model 2018** | **DeepViral 2020 (Seq)** | **DeepViral 2020 (Seq + Pheno + Function)** |
|---|---|---|---|---|---|
| **TR1-TS1** | ACC | 0.867 | 0.780 | 0.876 | 0.891 |
| | AUC | 0.912 | 0.886 | 0.934 | 0.907 |
| **TR2-TS2** | ACC | 0.840 | 0.780 | 0.792 | 0.903 |
| | AUC | 0.941 | 0.867 | 0.959 | 0.973 |
| **TR3-TS1** | ACC | 0.857 | 0.774 | 0.795 | NA |
| | AUC | 0.921 | 0.884 | 0.904 | NA |
| **TR4-TS2** | ACC | 0.900 | 0.817 | 0.824 | NA |
| | AUC | 0.956 | 0.890 | 0.966 | NA |

The two datasets containing human-virus interactions (TR1-TS1 and TR2-TS2) were applied to LSTM-PHV, Zhou’s model, the DeepViral that used amino acid sequences alone, and the DeepViral integrating three features. The other two datasets containing host-virus interactions (TR3-TS1 and TR4-TS2) were applied to the LSTM-PHV, Zhou's model and DeepViral. ACC and AUC correspond to accuracy and the area under the ROC curve, respectively. The performances were obtained from Table 1 in Liu-Wei’s paper (DeepViral).

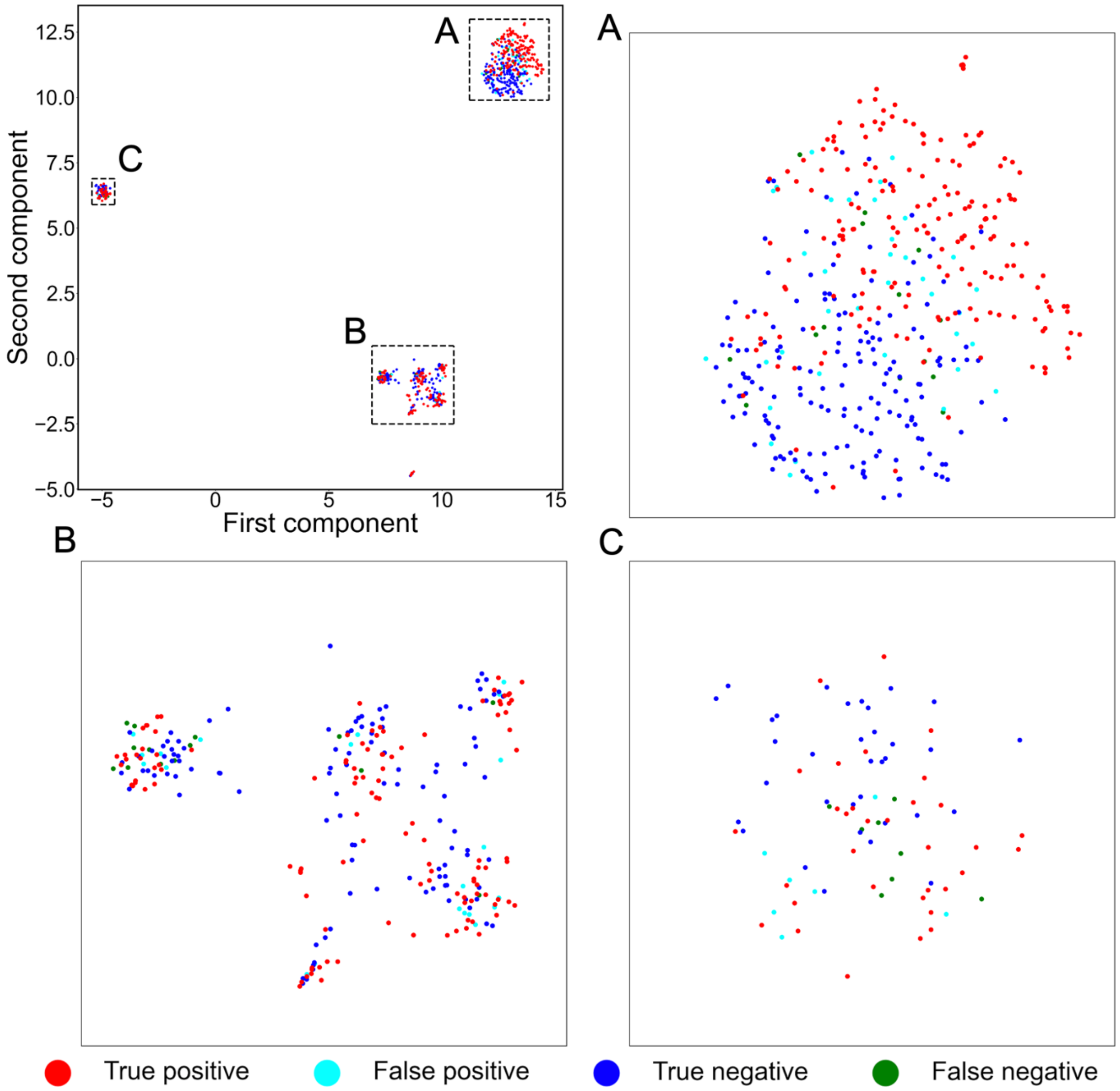


**Fig. S1.** UMAP of the concatenated vectors of TS1 predicted by the TR1-trained model. Three subspace (A, B, and C) in the main map were expanded in panels A, B, and C.

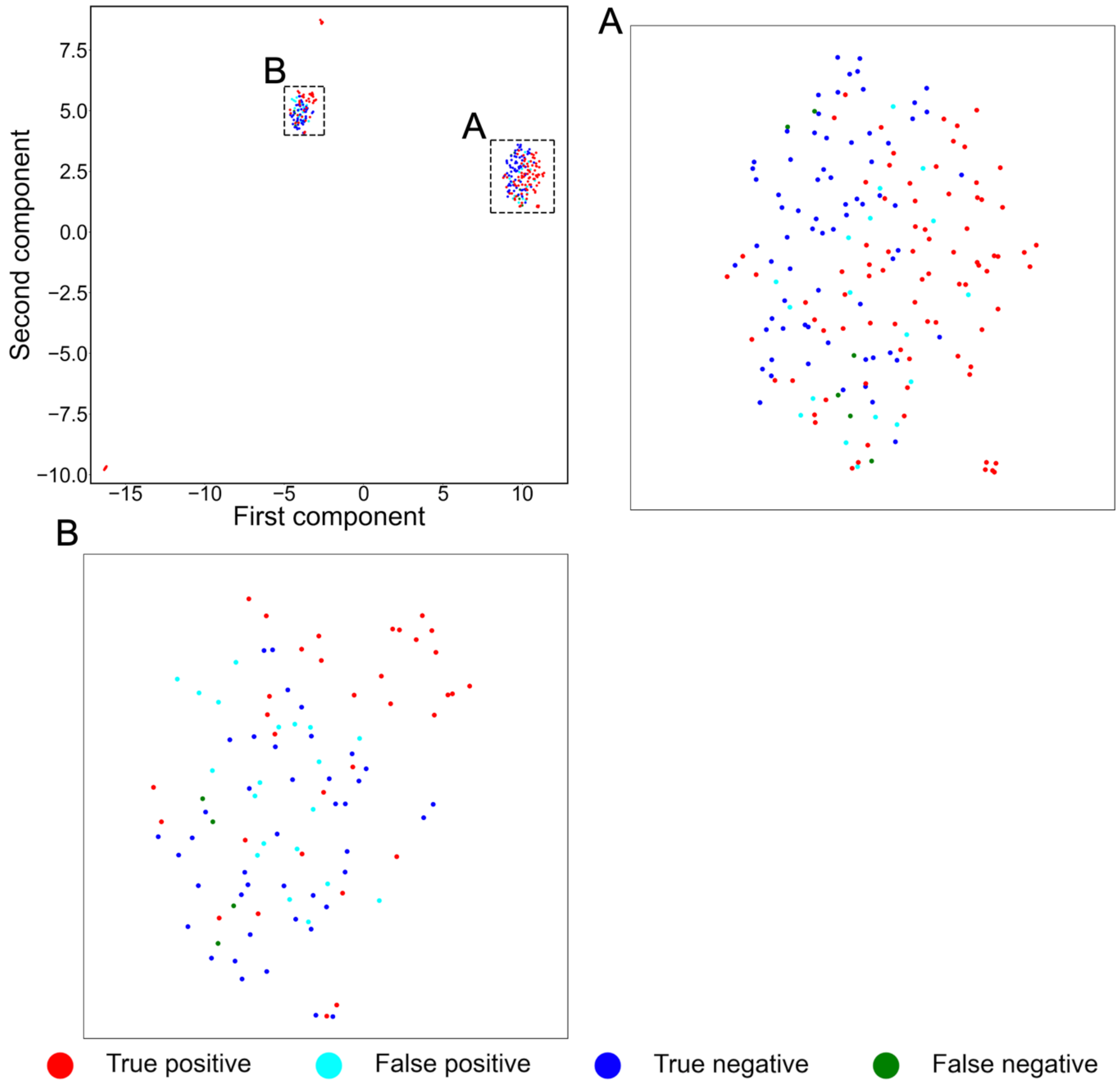


**Fig. S2.** UMAP of the concatenated vectors of TS2 predicted by the TR2-trained model. Three subspace (A, B, and C) in the main map were expanded in panels A, B, and C.

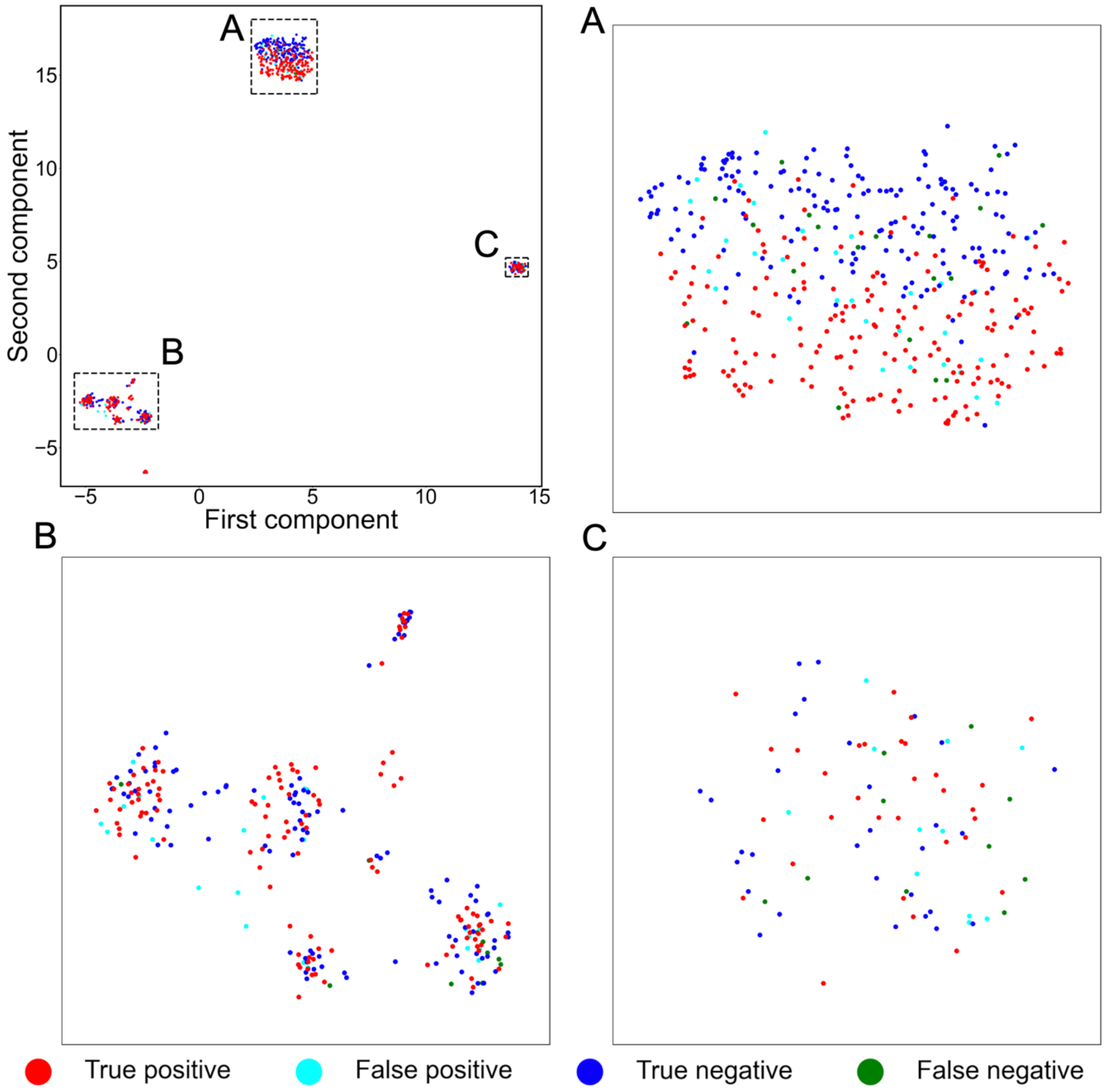


**Fig. S3.** UMAP of the concatenated vectors of TS1 predicted by the TR3-trained model. Three subspace (A, B, and C) in the main map were expanded in panels A, B, and C.

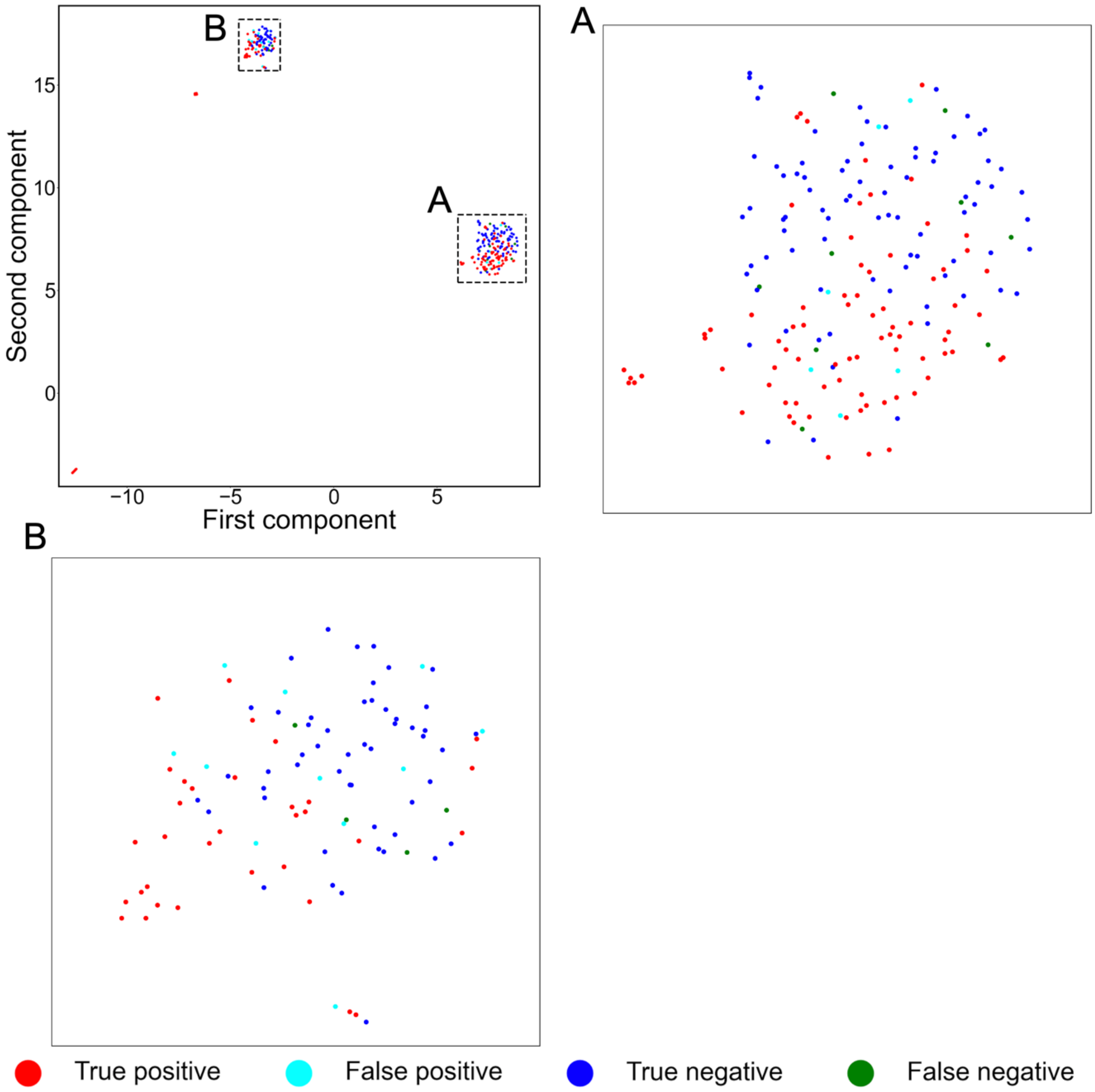


**Fig. S4.** UMAP of the concatenated vectors of TS2 predicted by the TR4-trained model. Three subspace (A, B, and C) in the main map were expanded in panels A, B, and C.
